# Addressing the Association Between Action Video Game Playing Experience and Visual Search in Naturalistic Multisensory scenes

**DOI:** 10.1101/2021.10.04.462750

**Authors:** Mohammad Hamzeloo, Daria Kvasova, Salvador Soto-Faraco

## Abstract

Prior studies investigating the effects of routine action video game playing have demonstrated improvements in a variety of cognitive processes, including improvements in attentional tasks. However, there is little evidence indicating that the cognitive benefits of playing action video games generalize from simplified unisensory stimuli to naturalistic multisensory scenes – a fundamental characteristic of natural, everyday life environments. In particular, it is unknown whether video game experience has an impact on cross-modal congruence effects when searching through such multisensory scenes. The present pre-registered compared the performance of action video game players (AVGPs) with non-video game players (NVGPs) on a visual search task for objects embedded in video clips of realistic scenes. We run two identical online experiments with gender-balanced samples, for a total N=130. Overall, the data replicated previous findings reporting search benefits when visual targets were accompanied by semantically congruent auditory events, compared to neutral or incongruent ones. However, according to the results, AVGPs did not consistently outperform NVGPs in the search task overall, nor they exploit multisensory cues more efficiently than NVGPs. Exploratory analyses with self-reported gender as a variable revealed a potential difference in response strategy between experienced male and female AVGPs when dealing with cross-modal cues. These findings suggest that the generalization of the advantage of AVG experience to realistic, cross-modal situations should be done with caution, and considering gender-related aspects.

---

The rapid development of smart technologies has led to an explosion in videogaming, which has brought video game playing and its consequences to the forefront of scientific attention. According to a growing body of evidence, video game playing is associated with improvements in a variety of cognitive and perceptual skills (Hisam et al., 2018; Oei & Patterson, 2015; Powers, Brooks, Aldrich, Palladino, & Alfieri, 2013; Strobach, Frensch, & Schubert, 2012). It has been suggested that at least some of the differences in basic cognitive and perceptual performance between action video game players (AVGPs) and non-video game players (NVGPs) are due to baseline individual differences (Boot et al., 2008), but a significant body of work in this domain has established that a causal relationship between playing action video games and improvements in attentional control (Bavelier & Green, 2019). It has been proposed that attentional control improvement due to playing action video games can be considered a training-related transfer between different perceptual and cognitive abilities (Ducrocq, Wilson, Vine, & Derakshan, 2016; Green, Gorman, & Bavelier, 2016). However, it remains largely unknown whether the improvement observed in AVGPs generalizes beyond simplified laboratory protocols to tasks involving the complexity of naturalistic, multisensory conditions of real-like scenarios. We set out to address this question.

In the literature, action video games are defined as a type of game that requires the processing of a large amount of visual information, presented rapidly over a wide field of view, and often requires the simultaneous tracking of multiple targets under high attentional demands (Green & Bavelier, 2006). These games include the so-called first- or third-person shooter or adventure games, such as *Medal of Honor* (Electronic Arts Inc.), *Call of Duty* (Activision Publishing Inc.) or *Grand Theft Auto* (Rockstar Games). In all cases, successful performance in these games requires extremely efficient attentional control.

Attentional control is a key mechanism of adaptive behavior and can be defined as our ability to stay focused on task-relevant information whilst resisting distraction from salient but irrelevant events, and being responsive to changes in the environment that require efficient re-orienting to new sources of relevant information (Engle, 2002). Thus, attentional control not only enables selective attention (focusing on spatial locations or objects that are goal-relevant while minimizing task-irrelevant information), but also the capacity to shift between selective and divided attention in order to consistently monitor changes in the environment (Corbetta & Shulman, 2002). One of the first scientific studies of visual attention in AVGPs, compared to NVGPs, reported that they had lower costs associated with targets appearing at low probability locations in a stimulus detection paradigm (Greenfield, DeWinstanley, Kilpatrick, & Kaye, 1994). Based on this finding, Greenfield et al. (1994) suggested that playing action video games may enhance the ability to allocate and divide attention, thereby improving visual search performance.

Over the last two decades, researchers have become interested in the visual search advantages of AVGPs. For example, AVGPs outperform NVGPs in traditional visual search tasks (Castel, Pratt, & Drummond, 2005; Hubert-Wallander, Green, Sugarman, & Bavelier, 2011), flanker/load tasks (Dye, Green, & Daphne Bavelier, 2009; Green & Bavelier, 2003; Green & Bavelier, 2006; Irons, Remington, & McLean, 2011; Xuemin & Bin, 2010), distraction-based tasks (Chisholm, Hickey, Theeuwes, & Kingstone, 2010; Rupp, McConnell, & Smither, 2016), and a change detection task (Clark, Fleck, & Mitroff, 2011; Durlach, Kring, & Bowens, 2009). These findings suggest that action video game experience enhances various aspects of top-down attention, and these effects have been found in both cross-sectional and in intervention studies (Bediou et al., 2018; Chisholm & Kingstone, 2015a; Schubert et al., 2015). Although it has often been proposed that these improvements are the result of changes in selective attention, more recently, enhancement in various additional aspects of top-down attention have been considered. For example, changes in attentional control or in the capacity to swiftly switch between attentional modes according to task demands (Bavelier & Green, 2019).

The vast majority of studies in this area have compared AVGP and NVGP performance on traditional laboratory protocols using simple visual stimuli. Furthermore, only a handful of studies have considered scenarios including sensory modalities other than vision. In a study measuring auditory decision making, AVGPs were found to be faster than NVGPs at indicating the ear to which the sound was presented, especially at low signal-to-noise ratios (Green, Pouget, & Bavelier, 2010). In a subsequent study using a demanding auditory discrimination task, AVGPs were able to detect auditory targets and to discriminate them from auditory non-target standards more accurately than NVGPs (Föcker, Mortazavi, Khoe, Hillyard, & Bavelier, 2019). In a cross-modal study, Donohue, Woldorff, and Mitroff (2010) compared AVGPs and NVGPs in an audiovisual simultaneity judgment task. The results showed that AVGPs were more accurate in discriminating whether pairs of simple visual and auditory stimuli were synchronous or slightly temporally offset. They also revealed an enhanced ability to judge cross-modal temporal order. Of note, Donohue et al.’s trials consisted of a small checker-board (5° by 5° visual stimulus) and a pure tone (1,200 Hz auditory stimulus), presented at varying onset asynchronies.

Despite the growing body of research on the benefits of action video game experience, few studies have investigated whether the performance benefits observed in simplified laboratory conditions can also be seen in more complex contexts with some of the features of real-world scenes. Taking a step towards more complex scenarios would be important to understand how the putative superiority of AVGPs in certain tasks transfers to naturalistic environments, which are complex, multisensory, and often semantically meaningful (Soto-Faraco et al., 2019; Kvasova, Stewart, & Soto-Faraco, 2021). In a study by Gaspar et al (2014), AVGPs and NVGPs were compared in a high-fidelity road crossing task in a virtual reality environment in two conditions: no distraction (single task) and a dual-task condition in which participants performed a continuous working memory task while crossing. They found that AVGPs and NVGPs performed similarly during road crossing in both conditions, and they concluded that the attentional benefits associated with AVG experience cannot transferred to complex road crossing performance involving multitasking. There is also some evidence that AVG experience is associated with better ‘real-life’ skills, such as higher psychomotor skills in laparoscopic surgery (Jalink et al., 2014; Lynch et al., 2010), alleviation of amblyopic symptoms (Li et al., 2015) or reading skills (Franceschini et al., 2013).

Recent studies on cross-modal interactions in attentional orienting have highlighted that in naturalistic scenarios, not only temporal and spatial congruence between stimuli across modalities plays a functional role in the control of attention, but also semantic correspondence can facilitate detection and recognition performance (Kvasova & Soto-Faraco, 2019; Roberts & Hall, 2008; Spagna, Mackie, & Fan, 2015). Cross-modal semantic facilitation has been demonstrated in a variety of tasks, including audiovisual matching task (Chen & Spence, 2010; Hein et al., 2007; Laurienti et al., 2003), visual awareness (Chen, Yeh, & Spence, 2011; Cox & Hong, 2015; Hsiao, Chen, Spence, & Yeh, 2012), spatial attention (List, Iordanescu, Grabowecky, & Suzuki, 2014; Mastroberardino, Santangelo, & Macaluso, 2015; Kvasova, Spence, & Soto-Faraco., 2023), and object search in naturalistic scenes (Kvasova, Garcia-Vernet, & Soto-Faraco, 2019; Pesquita, Brennan, Enns, & Soto-Faraco, 2013).

Here, we report the results of two studies using a visual search task in scenes that share some of the characteristics of realistic multisensory scenes, to answer two outstanding questions. First, whether AVGPs outperform NVGPs in a search task where they need to look for an object within complex, dynamic, multisensory scenes. Because naturalistic environments are multisensory, a second interrelated question we sought to answer is whether AVGPs can benefit more efficiently than NVGPs from cross-modal semantic congruence between visual events and their associated sounds when searching for objects.

In the present visual search task, which was similar to the task used in previous studies (Kvasova et al., 2019, 2021), targets consist of ordinary everyday life objects embedded in video clip fragments of complex naturalistic scenes. For example, participants were asked to search for a dog in a video-clip depicting a busy city street scene. A characteristic object sound (consistent with the search target, with a distractor object present in the scene, or unrelated to any objects in the scene) was presented at stimulus onset, mixed with ambient noise. Reaction times as well as visual search accuracy were measured. We must note that these stimuli are not strictly equivalent to the naturalistic real-life experience, but they do share many more of its characteristics than the artificial stimuli used in search tasks in most studies. For example, they are dynamic instead of static, they show familiar objects in contextualized interactions, they are embedded within semantically structured relations, and they are multisensory. In this respect, we surmise, the present study takes the issue of naturalistic generalization of the effects of gaming on attention performance one step further (see Matusz et al., 2019, for an example of how this incremental approach can help tackle the difficult problem of ecological validity). Likewise, there are also important differences with respect to a naturalistic situation, such as for example the 2D nature of the viewing display, the edited sounds, and the lack of spatial congruence of sounds to visual targets/distractors. The first difference is a simple matter of experimental convenience (the same reason that 2D displays have been used in a variety of other studies, e.g., Baldassano et al., 2018; Peelen & Kastner, 2011; Spiers & Maguire, 2007). The latter two differences relate to design features to control for otherwise important confounds in the interpretation of the results (see Methods).

We hypothesized that, (1) if AVGPs benefit from more efficient attentional control to direct their attention to target-relevant information while minimizing target-irrelevant information in multisensory environments, they would outperform NVGPs in the task of searching objects in videos of naturalistic scenes. We assumed that this advantage would be observed in faster reaction times and/or more accurate responses overall in the task. In addition, our second hypothesis was (2) if AVPGs have learned to use environmental multisensory cues more efficiently, then we expect a greater cross-modal advantage for AVGPs. That is, AVGPs would exploit cross-modal congruence to a greater extent than NVGPs. We expect this because previous studies have revealed that as AVGPs possess a greater attentional capacity to spill over task-irrelevant information in high-load perceptual scenarios (Dye, Green, & Bavelier, 2009; Green & Bavelier, 2003). This is consistent with the finding that NVGPs show a reduction in the magnitude of the flanker effect (Bavelier & Green, 2019). Please note that the present study addresses the association, but not the causality between AVGP and performance differences.

The hypotheses, methods and analysis pipelines for Experiment 1 were pre-registered prior to data analysis, with the exception of the Bayesian statistics which were added posteriori. The pre-registration is available at https://osf.io/sgy5a/. The results of the planned analyses are presented separately from the exploratory ones in the results section, below. Experiment 2 is a close replication of Experiment 1 and was not pre-registered specifically.

## Experiment 1

### Method

#### Participants

We used G*Power (Faul et al., 2009) to estimate the sample size for the study, using a repeated measures ANOVA with within-between interactions, a medium effect size f=0.25, and α level= 0.05. The total sample size for a statistical power of 0.95 (β = 0.95) was estimated to be N=54 (Tomczak, Tomczak, Kleka, & Lew, 2014). By considering inclusion and exclusion criteria, we enrolled 60 individuals using the online platform Prolific.co. Most of the participants were from European countries, with some others from South American countries, the USA, Canada, and one from South Africa. Based on their responses to a questionnaire asking about their video game playing habits in different genres during the past 12 months (Bavelier Lab Gaming Questionnaire), we had 30 AVGPs and 30 NVGPs. The mean age was 23.78 years and SD = 3.12. There was no significant age difference between the two groups (t = 0.115, df = 28, *p* = 0.91), and the two groups were gender balanced (15 males and 15 females within each group).

Otherwise, the general inclusion criteria were: (1) having a normal or corrected-to-normal vision and hearing, (2) having a good quality of internet connection and access to the experiment from a laptop or personal computer as the experiment could only be responded to with a keyboard, (3) being a proficient English language speaker, (4) having a false alarm rate in catch trials (trials in which the search target was not present) of less than 15%, and (5) having an average accuracy in the three target-present experimental conditions was more than 70%. Accuracy exclusion criteria were used to ensure data quality and to provide a sufficient number of trials for a reliable estimate of reaction times. Although these criteria may reduce the chances of detecting effects of accuracy, they leave room for variation in performance to be registered.

#### Design

The design includes two independent variables: Video-game experience (between subjects) and sound-target relation (within-subjects). The first variable, Video-game experience, was measured using the Bavelier Lab Gaming Questionnaire, version November 2019. Based on the questionnaire adapted from Green et al. (2017), we had two groups of participants according to their experience: AVGPs and NVGPs. An AVGP is a person who has played at least 5 hours per week of first/third-person shooter and/or action RPG/adventure genre of games in the past year. An NVGP is a person who has played no more than 1 hour per week of any video game genre of in the past year. This categorization of video game playing habits is based on Li, Polat, Makous, and Bavelier (2009).

The other variable in the design, sound-target relation, manipulated as a within-subjects independent factor, refers to the semantic relationship between the visual search target in the task and the object sounds on each trial. This variable had 3 levels: target-consistent sound, distractor-consistent sound, and neutral sounds. In the target-consistent condition, the sound was matched to the target object. In the distractor-consistent condition, the sound was matched to the distractor (a non-target object present in the scene), and in the neutral condition, the object sound was not semantically congruent with any object in the scene. In order to measure false alarm rates and to balance the response types (50% yes, 50% no) in the task, the task included additional catch trials in which targets were not present (hence, the correct response was ‘NO’). Within these catch trials, as in target-present trials, the object sound had 3 levels: target-consistent sound (here, the sound is consistent with the designated search object, which is not present in the video clips), distractor-consistent sounds, and neutral sounds. The dependent variable of our design was visual search object performance, measured by correct response times on target-present trials, and by *d’*, calculated from the hit rate in target-present conditions and the false alarm rate in target-absent (catch) trials.

#### Stimuli

The materials for the visual search task (target objects, sounds, and video clips) in naturalistic environments were selected from those used in Kvasova et al. (2019) (see Figure 1, for an example). There was a set of 156 different video clips that were extracted from films, TV shows, and advertisements, or recorded by Kvasova et al. from scenes of everyday life. The video clips were recorded/played in color, at 30 frames per second and a resolution of 1024 * 768 pixels. All video clips were cut into 2-second fragments with no fade-in or -out. Twelve videos were used for training trials, 72 videos were used for the three target-present trials (24 each; see Table 1), and another 72 videos were used for catch trials in which the search target object was not present (again, 24 videos in each condition). The original sound of the videos has been replaced by background noise, created by superimposing various everyday sounds that have been adapted to the context of the video. Each video clip in the target-present trials contained at least two visual objects with a familiar characteristic sound (e.g. musical instruments, animals, tools, …). One of these objects was designated as the search target and the other as the distractor. The selection of the target/distractor objects followed two criteria, which were checked by at least two judges during the selection of the videos: (1) They were visible but not a part of the main action in the scene. For instance, if a person is talking on the mobile phone as the main action in the scene, the mobile phone cannot be a target object. (2) Both target and distractor objects were present throughout the video clip.

**Figure 1.**
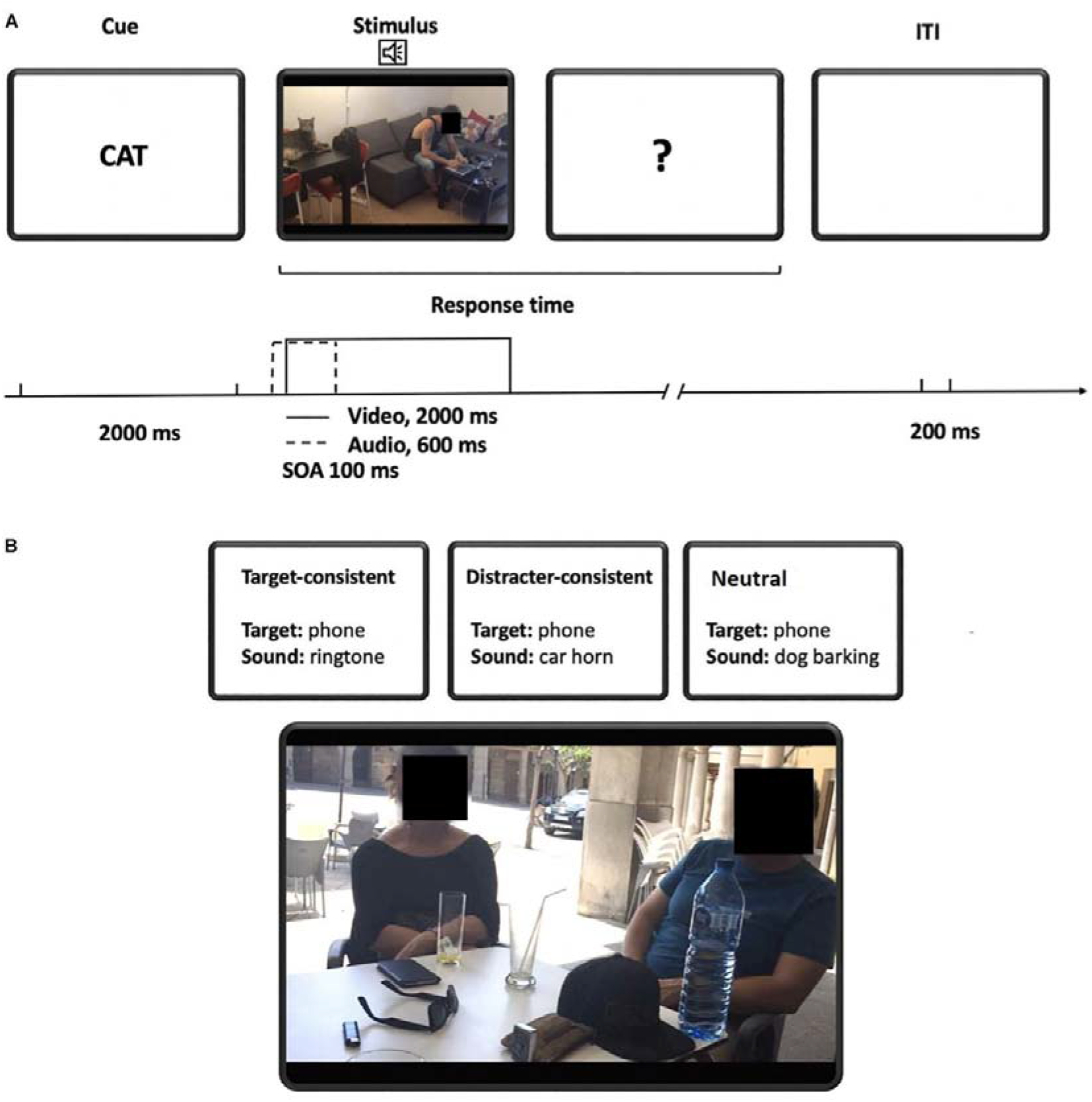
**(A)** Sequence of events in a typical experimental trial. The trial started with the presentation of the target word for 2000 ms, followed by the auditory cue and the video clip (see times in the figure). There was no time limit for the participant’s response. 200 ms after the participant’s response, a new target word was presented, beginning a new trial. **(B)** Example of sound conditions. In this stimulus example (shown with the snapshot of the video clip), the possible target (in target-present conditions) is the smartphone. In the target-consistent condition the sound corresponds to the target (e.g., a ring tone), in the distractor-consistent condition, the sound corresponds the distractor object (in this case, the car), and in the catch trial, the sound does not correspond to any object in the scene (e.g., the dog barking). The image is a frame from the video clip filmed by the research group.

**Table 1:** Distribution of trials and conditions.

|  | Sound Condition | Description | Number of trials |
| --- | --- | --- | --- |
| **Target-present trials** | Target-consistent | The target is present and the sound is congruent with the target object | 24 |
|  | Distractor-consistent | The target is present and the sound is congruent with a distractor object | 24 |
|  | Neutral | A target is present and the sound is not related to any object in the video | 24 |
| **Catch trials** | Target-consistent | The target is not present and the sound is congruent with the target word | 24 |
|  | Distractor-consistent | The target is not present and the sound is congruent with a distractor object | 24 |
|  | Neutral | The target is not present and the sound is not related to any object | 24 |
| **Total** |  |  | 144 |

The use of the two designated objects of each video as a target or distractor was counterbalanced across the experimental design to compensate for potential biases related to the specific objects. To reach this goal, we used each video for all three conditions in target-present trials. The three different experimental conditions were created by combining the video with target-consistent sound, distractor-consistent sound, and neutral sound while one of the two target objects was present on the screen before the video, to create six equivalent versions of a target-present trial (2 target objects * 3 conditions). In each video clip in the catch trials, there was at least one visual object with a characteristic sound, that was used as a distractor object. We created 3 equivalent versions of catch trials from each video, in the same way, so that each catch video appeared in the three conditions across different versions of the experiment. In the end, we had six different versions of the task of equal length including one version of target-present trials and one version of catch trials. Each version was presented to 10 participants with different random order of videos. Throughout the experiment, each video and possible target/distractor object appeared equally in each condition. This ensured that any variability between video clips could not explain differences between conditions. Characteristic sounds were semantically compatible with the target/distractor object, depending on the particular condition, and were presented centrally, providing no cues for object location. All the sounds were normalized at origin to have equivalent SPL and were presented for 600ms.

#### Procedure

We built the experiment using Builder under the Psychopy package (v. 2020.1.3) and ran it under the Pavlovia.org platform. Psychopy is also the only package with a reaction time precision of less than 4 ms online (Sauter, Draschkow, & Mack, 2020). We recruited participants through the Prolific.co. platform. Each participant was asked to read the consent form and confirm their agreement to participate in the experiment voluntarily. The study was approved by the UPF Institutional Committee for the Ethical Review of Projects (CIREP-UPF). They were able to exit the experiment at any moment by pressing the ‘Escape’ key in their keyboard. After consent was given, they filled a form of demographic data including age, gender (by selecting M for male, F for female, and O for other), and the video game questionnaire in the first part of the study. We continued running the first part of the study for 119 participants until we had 15 participants in 4 cells of gender x group in a stratified fashion. If they fell into each group category (AVGP or NVGP), they received an invitation to the second part of the study in less than an hour; if not, they were paid 0.5 £ as compensation for their participation. By clicking on the invitation link, the instructions for the main task appeared on the screen and they were asked to complete the task indoors with dim lighting and to turn off their phone to avoid distractions during the task. They were also asked to increase the volume of their device up to 80%, or to a level where they could easily hear the sounds. By pressing the spacebar, they entered a training block of 12 trials before the beginning of the experiment. In the training trials, feedback was provided on the participant’s response to ensure that they understood the task. No feedback was given during the experimental blocks.

Each trial began with a cue word designating the search target object which was displayed in the center of the screen for 2000ms. This cue was followed by the video clip, with the corresponding sound. Auditory cues (target-consistent, distractor-consistent, or neutral) began slightly ahead of the video, by 100ms, and lasted for 600ms. The video was shown for 2000ms. The participant judged whether or not the target object was present in the video clip. Participants were asked to put their left index finger on the "Y" key and their right index finger on the "N" key of their keyboard and to press the Y key if they found the target object in the video or press the N key if the target object was not presented. Participants were instructed to respond as quickly as possible, but there was no time limit for the trial response. A question mark was presented after the video offset, and until a response was made. There was a 200ms blank screen between trials.

When they finished the task, they were returned to the Prolific website where a successful submission appeared in their account. When their submission was approved by the researcher and they were paid 2.5£ as compensation for their participation. We collected 85 individual datasets, but 25 (17 NVGPs and 8 AVGPs) of them were excluded from the analyses due to the accuracy criteria (details provided in Table 2). We continued to run the experiment until we had 60 valid individual datasets to include in the analyses.

**Table 2:** Exclusion of individual subjects.

| Exclusion criteria | Group |  |
| --- | --- | --- |
|  | AVGPs | NVGPs |
| False alarm rate in catch trials (higher than 15%) | 3 | 8 |
| Average accuracy in experimental trials (less than 70%) | 4 | 3 |
| Both criteria | 1 | 6 |
| Total | 8 | 17 |

### Data analyses

#### Preprocessing

To eliminate other processes outside the interest of the study, such as quick lucky guesses, delayed responses due to subject’s inattention, or guesses based on the subject’s failure to reach a decision, we considered an outlier filter for RTs +/-2SD around the mean of each condition for each subject: neither RT nor accuracy data were analyzed for these outlier trials. After filtering incorrect trials from all target-present trials, and 3.89% of the data was filtered out using the above outlier filter criterion (137 out of 3,522 RTs).

#### Reaction times

In some of the within-subject analyses, normalized RTs were used where indicated. The data from neutral trials were used to normalize the data from the conditions of interest across participants, in order to reduce inter-individual differences or expectancy-related differences in performance (demand characteristics) that could be created by filling the Video Game Questionnaire in AVGP and NVGP participants, and to concentrate on the effects of interest. The normalization was performed according to Equation (1) for each subject (i) and condition of interest (j):

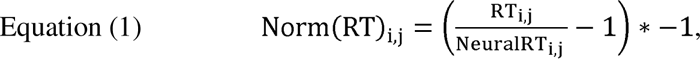

where *Norm*(*RT)_i,j_* is the normalized RT for subject *i* in condition *j*, (*RT)_i,j_* is the mean RT for subject *i* in condition *j*, and *Neutral(RT)_i,j_* is the mean RT in the neutral sound condition of subject *i*. The norm RTs equal to zero means that the RT for the condition of interest was equal to the RT for neutral condition which created a baseline. The positive norm RTs showed the cross=modal facilitation, and negative ones showed cross-modal distraction.

#### Performance data

Signal Detection was used to measure the precision of responses from the conditions of interest across participants by calculating *d’* according to Equation (2) for each subject (i) and condition of interest (j):

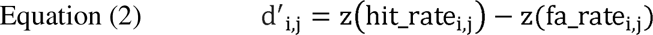

where *d’_i,j_* is the *d’* for subject i in condition j, *hit_rate _i,j_* is yes responses / total responses for subject i in condition j of target-present trials, *fa_rate _i,j_* is yes responses / total responses for subject i in condition j of catch trials, *z(hit_rate _i,j_)* and *z(fa_rate _i,j_)* are standardized z scores for *hit_rate _i,j_* and *fa_rate _i,j_* respectively. If hit_rate or fa_rate is equal to 0, it has been replaced by 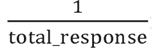 and if hit_rate or fa_rate is equal to 1, it has been replaced by 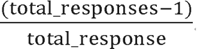.

In addition, *d’* scores from neutral trials were used to normalize the *d’* scores from the conditions of interest across participants according to Equation (3) for each subject (i) and condition of interest (j). The aim of the *d’* normalization was to investigate whether the congruency effects found in the conditions of interest, if any, were due to cross-modal facilitation in target detection, cross-modal distraction, or both.

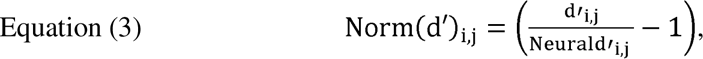

where *Norm*(*d’)_i,j_* is the normalized *d’* for subject *i* in condition *j*, (*d’)_i,j_* is the mean *d’* for subject *i* in condition *j*, and *Neutral(d’)_i,j_* is the mean *d’* in the neutral sound condition of subject *i*. The norm *d’* scores equal to zero means that the *d’* for the condition of interest was equal to the *d’* for neutral condition which created a baseline. The positive norm *d’* scores showed the cross-modal facilitation, and negative ones showed cross-modal distraction.

#### Bayesian analysis

To go beyond the limitations of standard Null Hypothesis Significance Testing (NHST) *p*-values, such as the inability to provide sufficient evidence for the absence of an effect (for a review see Kubsch, Stamer, Steiner, Neumann, & Parchmann, 2021), we conducted Bayesian analyses in parallel with each NHST in our study. Given that the goal of this study was to ascertain the presence or absence of the AVGP effect, we recruited Bayes Factor (BF) hypothesis testing for each effect to conclude whether the data are more likely under the research hypothesis (supporting the presence of the effect), the null hypothesis (supporting the absence of the effect), or whether there are ambiguous BFs indicating insufficient evidence to sufficiently support either hypothesis (for a more detailed discussion see, e.g., Dienes, 2014). All Bayesian analyses were implemented in the statistical software package JASP (JASPTeam, 2021).

## Results

### Results of the pre-registered analyses

The descriptive statistics for Experiment 1 are shown in Table 3. To evaluate our first hypothesis, whether the advantage of action video game experience can be transferred to search in complex, multisensory scenes, we addressed whether there is a superiority of AVPGs in speed and/or precision (*d*’). To test this, we used the inter-subject averages per group (AVPG, NVGP) pooled across all other conditions and performed a one-tailed t-test on RTs of correct responses to target-present trials, and on *d’* scores. The result showed significant differences between groups on mean RTs (t_(58)_ = 2.61, *p* < 0.01, *Cohen’s d* = 0.68, BF_10_ = 8.43) (Figure 2A), whereas no such difference occurred in the *d’* total (t_(58)_ = 0.63, *p* = 0.265, *Cohen’s d* = 0.16, BF_10_ = 0.45) (Figure 2B). These results confirmed our first hypothesis, suggesting that AVGPs responded faster, albeit with similar precision than NVGPs. In particular, the average advantage of AVPGs was of 220ms, a 14% variation over the total RT, which can be considered as moderate evidence with a medium effect size. Note that the lack of group differences in precision may be related to the accuracy-based exclusion criteria, which were introduced precisely to render latency estimations interpretable.

**Figure 2.**
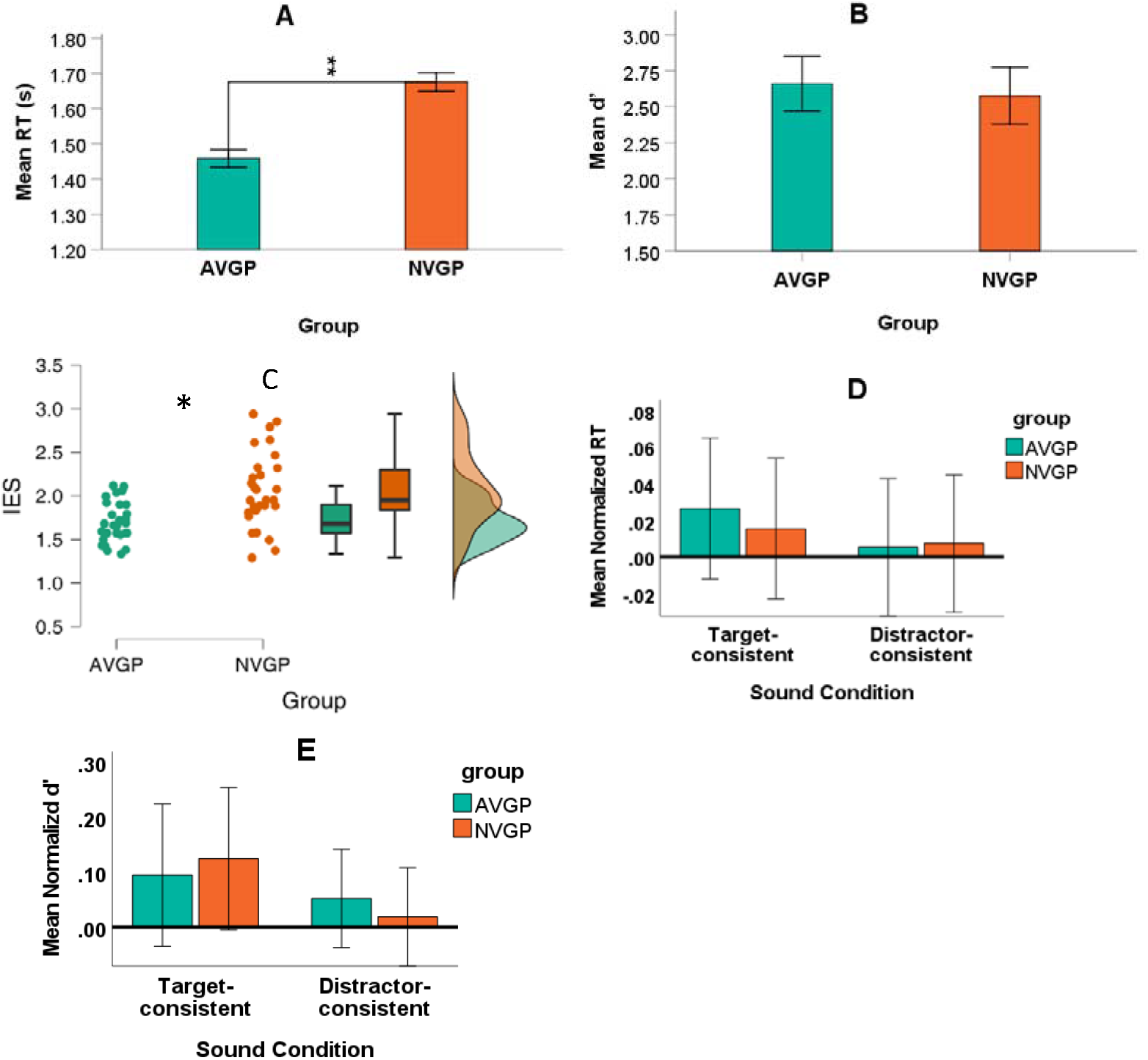
**(A)** Visual search Mean reaction times for correct responses to target-present trials, plotted separately for AVGPs and NVGPs. (B) Total visual search accuracy (*d’*) plotted for AVGPs and NVGPs. **(C)** The inverse efficiency scores on a raincloud plot with means and confidence interval (95%) for the two groups. **(D)** Normalized reaction times in the target-consistent and distractor-consistent conditions plotted for AVGPs and NVGPs. **(E)** Normalized *d’* scores in the target-consistent and distractor-consistent conditions plotted for AVGPs and NVGPs. In all plots, error bars indicate the standard error of the mean. Significant differences are indicated by asterisks (*p-value<0.05; **p-value<0.01).

**Table 3:** Descriptive statistics (means and standard deviations) for Experiment 1.

| Variable | AVGP (n=30) |  |  | NVGP (n=30) |  |  |
| --- | --- | --- | --- | --- | --- | --- |
|  | Male | Female | Total | Male | Female | Total |
| Age | 24.67(2.92) | 23.93(2.96) | 24.30(2.91) | 25.67(4.05) | 23.80(3.36) | 24.23(3.68) |
| RTs for TC | 1.40(.12) | 1.42(.28) | 1.41(.21) | 1.73(.40) | 1.58(.24) | 1.65(.33) |
| RTs for DC | 1.46(.13) | 1.45(.46) | 1.45(.33) | 1.70(.40) | 1.65(.32) | 1.68(.36) |
| RTs for NC | 1.40(.13) | 1.53(.44) | 1.47(.33) | 1.74(.39) | 1.63(.22) | 1.69(.32) |
| RTs for TT | 1.42(.44) | 1.51(.61) | 1.46(.53) | 1.73(.59) | 1.62(.53) | 1.68(.56) |
| Norm RTs for TC | 0.00(.10) | 0.05(.11) | 0.03(.11) | 0.00(.13) | 0.03(.08) | 0.02(.10) |
| Norm RTs for DC | -0.04(.10) | 0.06(.10) | 0.01(.11) | 0.02(.09) | -0.01(.10) | 0.01(.10) |
| $D'$ for TC | 2.52(.52) | 2.68(.33) | 2.60(.43) | 2.72(.51) | 2.51(.47) | 2.61(.49) |
| $D'$ for DC | 2.79(.40) | 2.36(.40) | 2.57(.45) | 2.59(.56) | 2.25(.60) | 2.42(.60) |
| $D'$ for NC | 2.71(.53) | 2.33(.59) | 2.52(.58) | 2.72(.48) | 2.16(.40) | 2.44(.52) |
| $D'$ Total | 2.8(.55) | 2.52(.44) | 2.66(.51) | 2.77(.59) | 2.38(.39) | 2.58(.53) |
| Norm $d'$ for TC | -0.05(.20) | 0.24(.45) | 0.09(.37) | -0.03(.27) | 0.22(.38) | 0.12(.34) |
| Norm $d'$ for DC | 0.05(.17) | 0.06(.25) | 0.05(.21) | -0.02(.30) | 0.05(.26) | 0.02(.28) |

TC= Target consistent condition; DC= Distractor consistent condition; NC= Neutral condition; TT= Target-present Trials.

A potential concern with the pre-registered protocol was that we set performance exclusion criteria across the board, which may have tended to select high performing NVGPs, hence equating performance (that is, *d’* scores) across groups. Because if this were the case, it would in fact go against the predicted results of group differences in RTs, the significant group difference in RTs reinforces the conclusion. Nevertheless, we decided to evaluate our hypothesis further, to compensate for potential speed-accuracy trade-offs between the two groups. In an exploratory analysis we analyzed Inverse Efficiency Scores (IES) calculated based on Equation (3) to combine speed and accuracy in a single metric. The result of a one-tailed t-test on IES showed a significant difference between groups (t_(58)_ = 3.01, *p* < 0.005, *Cohen’s d* = 0.78, BF_10_ = 9.92) (Figure 2C), in the expected direction, and with a slightly larger effect size than the RT effect. Finally, we also repeated the RT and *d’* analyses including all N=85 subjects tested (that is, including those that had been initially filtered out given the pre-registered exclusion criteria), and report them in the Supplementary Materials. In short, the data were more variable (given the inclusion of low-quality datasets) but the statistical tests provided equivalent results in RT, and a small but significant *d’* effect in the expected direction).

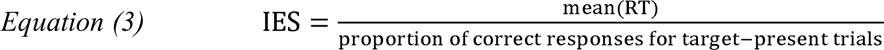

To evaluate our second hypothesis, whether AVGPs can benefit more from congruent cross-modal cues and/or are more resistant to cross-modal distractors, we entered the normalized RTs into a repeated-measures analysis of variance (ANOVA) with the between-participants factor of group (AVGPs vs. NVGPs) and the within-participants factor sound condition (target-congruent and distractor congruent, normalized by RTs in the neutral condition). Please note that to address this test, it is important to compare relative effects (the cross-modal advantage) between the groups, as AVGPs and NVGP may have different baseline performance (as was indeed shown in the analyses above). The use of normalization would allow us to directly test for potential differences in cross-modal advantage, without the confounding of the overall differences between groups. Neither the interaction of group* sound condition (F_(1, 58)_ = 0.218, *p* = 0.64, η_p_^2^=0.001, BF_10_ = 0.06) (Figure 2D), nor the main effect of sound condition was significant (F_(1, 58)_ = 1.10, *p* = 0.29, η_p_^2^=0.019, BF_10_ = 0.23). The effect of group also showed a non-significant result (F_(1, 58)_ = 0.04, *p* = 0.85, η_p_^2^=0.001, BF_10_ = 0.06), unsurprisingly, given the normalization.

We ran another repeated-measures ANOVA on normalized *d’* scores with the same factors and, similar to the RTs, neither the interaction of group* sound condition (F_(1, 58)_ = 0.56, *p* = 0.46, η_p_^2^=0.010, BF_10_ = 0.02), nor the main effect of sound conditions was significant (F_(1, 58)_ = 3.09, *p* = 0.08, η_p_^2^=0.05, BF_10_ = 0.14) (Figure 2E). Group also showed a non-significant result (F_(1, 58)_ = 0.01, *p* = 0.98, η_p_^2^=0.000, BF_10_ = 0.06). The analyses then did not confirm our second hypothesis, and in fact Bayesian analyses provided strong evidence for H_0_, suggesting that AVGPs do not seem to benefit more from congruent cross-modal cues than NVGPs in speed or accuracy. They also seem not to be less distracted by incongruent cross-modal cues than NVGPs.

## Experiment 2

The results of Experiment 1 showed that AVGPs were faster than NVGPs in the visual search task, without any accuracy loss (if anything, *d’* scores went in the same direction as RTs). The conclusion supported by this first study, with regard to overall search performance, is that it extrapolates AVGPs search advantage over NVGPs to more complex, dynamic multisensory scenarios. However, we did not find any evidence that AVGPs profited from cross-modal congruence any more than NVGPs did, a null finding. Given this pattern of findings, we decided to run a replication experiment. First, in order to confirm the lack of group differences regarding crossmodal advantage. Second, and while we were at it, to address one potential concern we were made aware of: we had not control over the size and quality of the monitors that participants used to run the task. It can be assumed that AVGPs tend to have larger, better-quality screens with higher resolution. Please note despite we did not control (or record) participants’ monitor resolution and refresh rate, our online protocol included the calibration of the size of the visual stimuli (video clips) and frame rates were equated across participants. We decided replicating Experiment 1 while controlling for the above variables.

In addition, in Experiment 1 response accuracy of the two groups did not differ significantly. It could be criticized that the equivalence in response accuracy could be due to performance-based participant selection criteria. The pre-set exclusion criteria were intended to insure data quality (helped to filter out participants with low attention or motivation), bringing performance within a measurable range and making RTs interpretable, whilst leaving plenty of room for these parameters to vary as a function of the factors of interest. Yet, one could argue our pre-set criteria where too strict, therefore masking an actual accuracy effect, in addition to the effect in response time. To further clarify this issue, we ran the analyses for Experiment 2 twice, once with the same selection criteria reported in the main text to replicate the above results, and once without the selection criteria reported in the Supplementary materials.

### Methods

The design, stimuli, and procedure of Experiment 2 were the same as Experiment 1 except for the following aspects. First, we used a new version of the Bavelier Lab Gaming Questionnaire (November, 2022). The new version incorporates two questions for each type of game to specify whether the participant played more than half of the time on a small screen, meaning 12 inches or smaller, and/or using a touchscreen. These two questions were used to avoid including individuals who play games on smartphones and tablets. We also recorded the resolution, and size of the monitor people used to perform the task.

#### Participants

To calculate the sample size, we used the effect size (Cohen’s *d* = 0.68) obtained from the analysis of the first hypothesis in Experiment 1. Considering a t-test for comparing two independent means, αlevel= 0.05 and β = 0.95 in G*Power software (Faul et al., 2009), the total sample size was calculated to be 96. After administering the video game questionnaire, we sent 100 invitations to those who fell into each group category (AVGP or NVGP) based on their gaming habits in the past 12 months. We collected 96 individual records. Considering the accuracy criteria (hit rate above 85% for catch trials and 70% for target-present trials), we excluded 26 data (14 AVGPs and 12 NVGPs). The final sample was 70 data including 36 AVGPs (16 female, M = 24.50, SD = 4.03) and 34 NVGPs (18 female, M = 28.06, SD = 4.36). The analyses with all 96 data (50 AVGPs and 46 NVGPs) can be found in the supplementary materials.

## Results

Before conducting the analysis for the main hypotheses, we compared the two groups according to the size of the monitors they used to run the experiment. As expected, AVGPs used larger monitors (M = 19.39**"**, SD = 5.52) than NVGPs (M = 17.64**"**, SD = 3.64) (t_(68)_ = 2.05, *p* < 0.05). However, screen size is not a good indicator of the quality. As we recorded the participants’ screen resolution in pixels and size in inches, we calculated the pixels per inch (PPI) as a measure of pixel density using Equation (4). The results showed that there was no significant difference between the two groups in terms of PPI (t_(68)_ = 0.65, *p* = 0.26), meaning that the quality of the screens in terms of the number of pixels per 1-inch line was the same for both groups. Therefore, it seems that the size of the screen could not affect the results of the visual search task (please note that the size of the stimulus was equated as part of the online protocol).

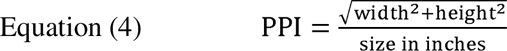

After filtering RT outliers from all target-present trials, 1.17% of the data was filtered out using the outlier filter criterion (RTs +/-2SD around the mean of each condition for each subject) (81 out of 6907 RTs).

The descriptive statistics for Experiment 2 are shown in Table 4. We performed two one-tailed t-tests on RTs of correct responses to all target-present trials and on *d’* scores to assess our first hypothesis. The results showed that there wasn’t any significant difference between the two groups in terms of RTs (t_(68)_ = 0.59, *p* = 0.72, *Cohen’s d* = 0.14, BF_10_ = 0.29) and *d’* scores (t_(68)_ = 0.05, *p* = 0.52, *Cohen’s d* = 0.01, BF_10_ = 0.25). Although the result of *d’* scores is consistent with the result of Experiment 1, the RTs outcome is inconsistent with Experiment 1. The result of a one-tailed t-test on IES showed a non-significant difference between the groups too (t_(68)_ = 0.74, *p* = 0.77, *Cohen’s d* = 0.18, BF_10_ = 0.31) (Figure 2C). These findings do not support our first hypothesis, as they suggest that AVGPs did not outperform NVGPs in the task of searching for objects in videos of realistic, dynamic complex scenes.

**Table 4:**
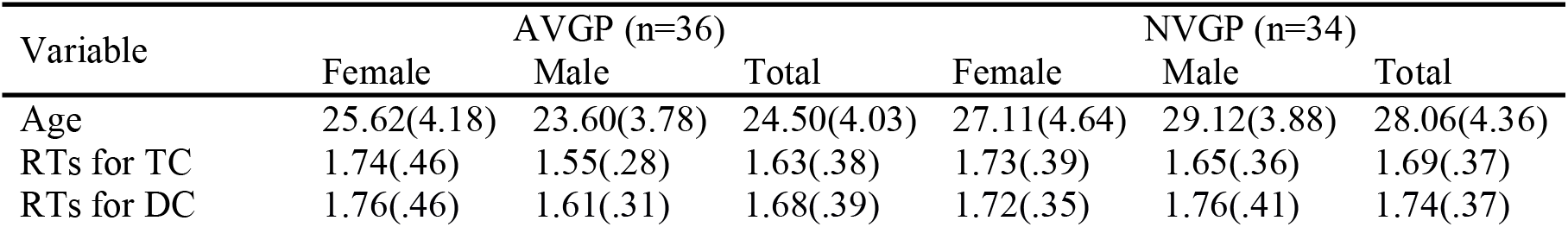
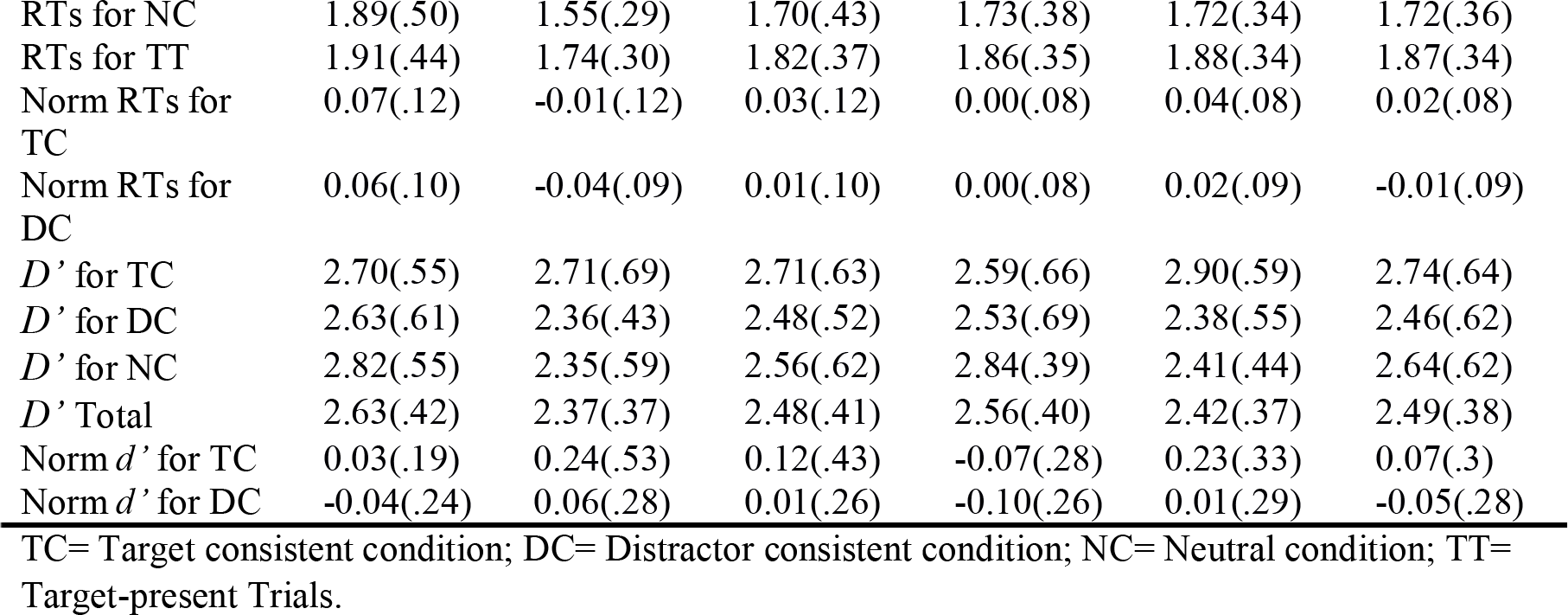
Descriptive statistics (means and standard deviations) for Experiment 2.

To evaluate our second hypothesis, we entered normalized RTs into a repeated-measures analysis of variance (ANOVA) with group as a between-subjects factor (AVGPs vs. NVGPs) and sound condition as a within-subjects factor (target congruent and distractor congruent, normalized by RTs in the neutral condition). The result of the analysis for the between-subjects factor (group) was not significant (F_(1, 68)_ = 0.35, *p* = 0.55, η_p_^2^=0.005, BF_10_ = 0.30). Sound condition had a significant main effect (F_(1, 68)_ = 5.77, *p* < 0.05, η_p_^2^=0.02, BF_10_ = 3.99), while the interaction of group*sound condition was not significant (F_(1, 68)_ = 0.17, *p* = 0.68, η_p_^2^=0.002, BF_10_ = 0.37). We ran another repeated-measures ANOVA on normalized *d’* scores with the same factors. The test of between-subjects factor (group) was not significant (F_(1, 68)_ = 0.58, *p* = 0.45, η_p_^2^=0.009, BF_10_ = 0.26). The sound condition had a significant main effect (F_(1, 68)_ = 8.94, *p* < 0.01, η_p_^2^=0.121, BF_10_ = 4.27), while the result of the interaction between the group and sound condition was not significant (F_(1, 68)_ = 0.02, *p* = 0.88, η_p_^2^=0.000, BF_10_ = 0.26). Figures 3D and 3E plotted the facilitation effect of target consistent sound in both reaction time and the accuracy of responses for the both groups, whereas distractor congruent sound did not show neither the facilitation nor distraction effect. These results, consistent with the outcome of Experiment 1, do not support our second hypothesis and suggested that AVGPs and NVGPs benefited equally from cross-modal congruency in speed or accuracy. They also seem not to be distracted by incongruent cross-modal cues less than NVGPs.

**Figure 3.**
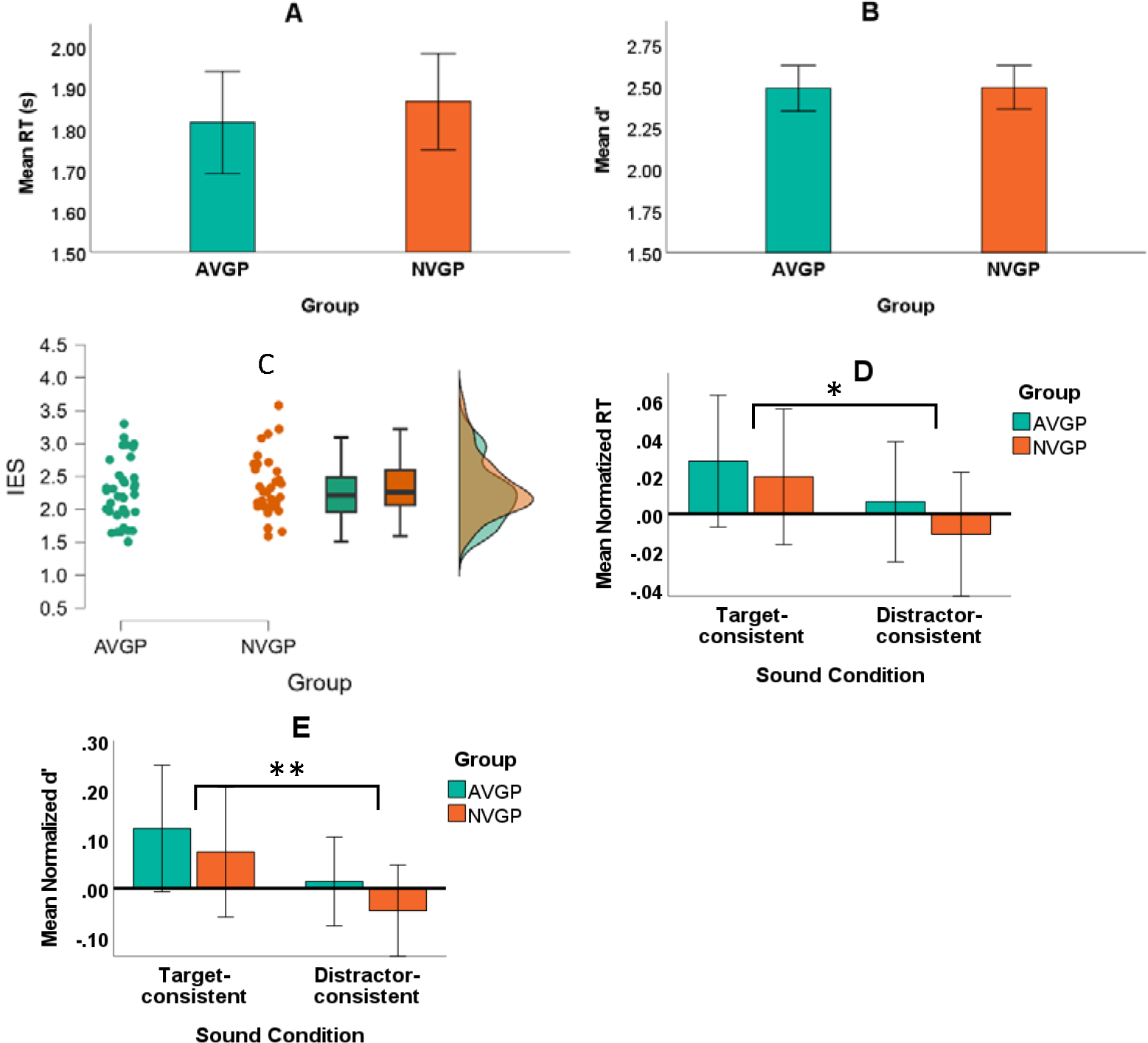
**(A)** Visual search Mean reaction times for correct responses to target-present trials, plotted separately for AVGPs and NVGPs. (B) Total visual search accuracy (*d’*) plotted for AVGPs and NVGPs. **(C)** The inverse efficiency scores on a raincloud plot with means and confidence interval (95%) for the two groups. **(D)** Normalized reaction times in the target-consistent and distractor-consistent conditions plotted for AVGPs and NVGPs. **(E)** Normalized *d’* scores in the target-consistent and distractor-consistent conditions plotted for AVGPs and NVGPs. In all plots, error bars indicate the standard error of the mean. Significant differences are indicated by asterisks (*p-value<0.05; **p-value<0.01).

**Figure 4.**
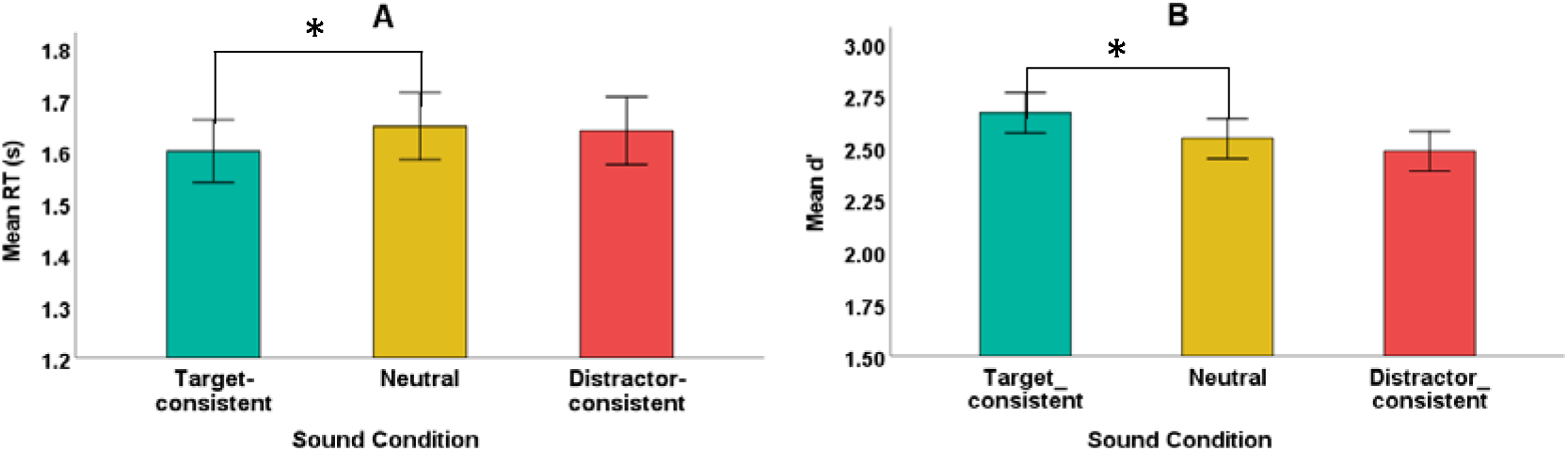
**(A)** Average reaction times and **(B)** *D’* scores in each of the three experimental sound conditions are plotted separately. Error bars indicate the standard error of the mean. Significant differences are indicated by asterisks (*p-value<0.05).

## Exploratory analyses with pooled datasets

In addition to the hypothesis-driven analyses described above, we pooled the two datasets from Experiments 1 and 2 together and performed two further exploratory analyses which were not directly addressed in our initial hypotheses:

***(1) Overall cross-modal effects.*** We addressed a comparison between the neutral and target-consistent condition, and between the neutral and distractor consistent condition across the board. This analysis shall be informative as to whether overall, congruent cross-modal cues benefit performance, incongruent cross-modal cues hinder performance, or both. Also, this analysis will help situate the present data with respect to previous studies finding cross-modal benefits in visual search. We ran two one-sided paired samples t-tests to compare mean RTs in the target-consistent condition and the distractor consistent condition to the mean RTs of the neutral condition. The results showed a significant RT advantage of target-consistent condition over neutral condition with strong evidential value (t_(129)_ = 3.09, *p* < 0.01, *Cohen’s d*=0.27, BF_10_ = 18.105). The difference between mean RTs of distractor-consistent condition and neutral condition was, however, not significant, with substantial evidence for lack of an effect (t_(129)_ = 0.623, *p* = 0.267, *Cohen’s d*=0.05, BF_10_ = 0.172) (Figure 3A). We ran two more one-sided paired samples t-tests, this time to compare *d’* of target-consistent condition and distractor consistent condition to the neutral condition. The results showed a significant higher *d’* in the target-consistent condition compared to the neutral condition but with low evidential value (t_(129)_ = 1.872, *p* = 0.032, *Cohen’s d* = 0.16, BF_10_ = 1.026) (Figure 3B). Similar to the RT outcome, there was no significant difference between *d’* of distractor-consistent condition and neutral condition (t_(129)_ = 1.101, *p* = 0.136, *Cohen’s d* = 0.097, BF_10_ = 0.303). Altogether, these results indicate that congruent cross-modal cues tend to benefit performance, but incongruent cross-modal cues do not hinder performance, with respect to neutral conditions. These results replicate recent findings from Kvasova et al. (2019) using the same search task on naturalistic scenes, and are in line with several previous cross-modal findings using more traditional laboratory tasks (Iordanescu, Guzman-Martinez, Grabowecky, & Suzuki, 2008; Knoeferle, Knoeferle, Velasco, & Spence, 2016; Kvasova, Spence, & Soto-Faraco., 2023; Laurienti et al., 2003).
***(2) Pooled datasets analyses and gender-dependent effects*.** We run analyses of the main hypotheses using the pooled datasets from Experiment 1 and 2. Given that we collected data from a gender-balanced sample in the both experiments for both AVGP and NVGP groups, we decided to include exploratory analyses of the two main hypotheses, considering gender as a between-subjects variable. Please note that initially gender was balanced in order to obtain measures from a more representative sample and prevent selection bias, but not to undergo any systematic investigation of the effects of gender. The participant selection by the online platform included male/female/others, and corresponds to the gender chosen by participants at the moment of running the experiment. We only had one data point where gender was selected as ’other’, which was excluded from the analyses due to independent accuracy criteria. For these reasons, we note here that the outcomes of these analyses must be carefully interpreted, given that the study was underpowered in terms of investigating interactions with the gender variable.

Regarding the first hypothesis (overall advantage of AVGPs over NVGPs), the result of a group (AVGP/NVGP) by gender (male/female) between-subject ANOVA on mean RTs revealed neither a group*gender interaction nor a main effect of gender (F_(1, 126)_ = 1.43, *p* = 0.23, η_p_^2^=0.011, BF_10_ = 0.15; F_(1, 126)_ = 0.19, *p* = 0.66, η_p_^2^=0.001, BF_10_ = 0.17 respectively). The main effect of group here also was not significant (F_(1, 126)_ = 3.50, *p* = 0.064, η_p_^2^=0.027, BF_10_ = 0.69). So, the outcome of the analysis using the pooled dataset did not give support to the hypothesis of AVPG advantage in search, in this task. We ran another group*gender between-subject ANOVA on *d’* scores. The result indicated no significant effects: the group*gender interaction (F_(1, 126)_ = 0.70, *p* = 0.40, η_p_^2^=0.006, BF_10_ = 0.04), the main effect of gender (F_(1, 126)_ = 0.295, *p* = 0.59, η_p_^2^=0.002, BF_10_ = 0.15), and the main effect of group (F _126)_ = 0,17, *p* = 0.68, η_p_^2^=0.001, BF_10_= 0.14). Collectively, the RT and *d’* outcomes using the pooled gender-balanced dataset of 170 participants reveal that AVGPs do not show any advantage over NVGPs in our visual search task, with certain evidential value for the null hypothesis.

Regarding the second hypothesis (differential cross-modal advantage for AVGPs), we entered gender and group as between-subjects factors and sound condition as a within participant variable in a repeated-measures ANOVA on normalized RTs. The test of within-subjects effects returned a significant main effect for sound condition (F_(1, 126)_ = 5.67, *p* < 0.05, η_p_^2^=0.04, BF_10_ = 1.35), but none of the interactions between sound condition*group (F_(1, 126)_ = 0.008, *p* = 0.93, η_p_^2^=0.00, BF_10_ = 0.24), sound condition*gender (F_(1,126)_= 1.41, *p* = 0.24, η_p_^2^=0.011, BF_10_ = 0.49), and sound condition*group*gender (F_(1, 126)_ = 0.51, *p* = 0.48, η_p_^2^=0.004, BF_10_ = 0.13) was significant. The test of between-subjects factors showed a significant main effect of gender (F_(1, 126)_ = 6.58, *p* < 0.05, η_p_^2^=0.05, BF_10_= 7.67) and significant interaction between gender*group (F_(1, 126)_ = 9.55, *p* < 0.01, η_p_^2^=0.07, BF_10_ = 6.71), but the main effect of group was not significant (F_(1, 126)_ = 0.57, *p* = 0.45, η_p_^2^=0.05, BF_10_ = 1.55). Post-hoc analyses with Bonferroni correction showed that there was a significant difference between male and female in the AVGPs group (M_(difference)_ = 0.1, *p* < 0.01) in normalized RT for distractor-consistent condition, and a marginally significant difference on normalized RT for target-consistent condition (M_(difference)_ = 0.067, *p* = 0.052). These results indicated that female AVGPs benefited more from cross-modal congruency in their RTs and were not distracted by cross-modal incongruency, whereas male AVGPs showed more distraction, that is their RTs showed down on distractor-consistent trials.

We conducted another repeated-measures ANOVA on normalized *d’* scores by entering gender and group as between-subjects factors and sound condition as a within-subjects factor. The test of within-subjects effects returned a significant main effect for sound condition (F_(1, 126)_ = 11.63, *p* < 0.01, η_p_^2^=0.08, BF_10_ = 0.86) but none of the interactions between sound condition*group (F_(1, 126)_ = 0.37, *p* = 0.54, η_p_^2^=0.003, BF_10_ = 0.016), sound condition*gender (F_(1, 126)_ = 0.007, *p* = 0.93, η_p_^2^=0.000, BF_10_ = 0.019), and sound condition*group*gender (F_(1, 126)_ = 0.55, *p* = 0.46, η_p_^2^=0.004, BF_10_= 0.001) was significant. The test of between-subjects effects showed that neither the main effect of gender (F_(1, 126)_ = 0.69, *p* = 0.41, η_p_^2^=0.005, BF_10_ = 0.071) and group (F_(1, 126)_ = 0.32, *p* = 0.57, η_p_^2^=0.003, BF_10_ = 0.062), nor the interaction between gender*group was significant (F_(1, 126)_ = 0.05, *p* = 0.82, η_p_^2^=0.001, BF_10_ = 0.017) was significant. These results indicated that AVGPs did not show an advantage of cross-modal congruency compared to NVGPs in their response accuracy.

## General Discussion

The first aim of the present study was to assess whether the visual search advantage demonstrated by AVGPs over NVGPs with simple stimuli in typical laboratory attention tasks would generalize to a multisensory search display with naturalistic scenes. Using a visual search task in video clips of naturalistic scenes, with audiovisual congruent/incongruent cross-modal cues, Experiment 1 initially suggested that the AVGPs advantage extrapolates to naturalistic scenarios in terms of faster reaction times without speed-accuracy trade-off. However, this advantage of the AVGP was not replicated in Experiment 2 neither when pooling the two datasets in exploratory analyses. Given that Experiment 2 was run to better control for potential differences in the screen size than in Experiment 1, one would have to conclude that these differences may have been at the root of AVGP vs. NVGP differences seen in Experiment 1. Could this be a very small effect with inconsistent outcomes given the sample size? We believe that the pooled datasets analyses (N=130), which returned null differences with evidential value for absence of an effect based on Bayesian statistics may alleviate this potential concern. Although it may be criticized that one limitation of the current study in testing the transfer of the AVGP advantage to the visual search task in naturalistic displays was due to the participants exclusion criteria, our results remained unchanged even when including data from all subjects tested regardless of the performance criteria (see Supplementary Materials). These results, therefore, cast some reasonable doubt on the idea that the advantage from extensive action video game experience may transfer to search in naturalistic scenes under multisensory conditions.

The literature on transferring the advantage of action video game experience to cross-modal and complex tasks is ambiguous. While some previous studies on visual and multisensory tasks have showed an advantages of AVGP experience (Chisholm & Kingstone, 2012, 2015a; Di Luzio et al. 2021; Donohue et al., 2010; Stewart, Martinez, Perdew, Green, & Moore, 2019), others have failed to show an advantage in cross-modal and complex tasks (Delmas et al., 2022; Gaspar et al., 2014; Prevratil et al., 2022; Stewart et al., 2020). These results suggest that the AVGP advantage may be modality-specific, i.e. the advantage can be transferred to visually familiar or game-like tasks, but cannot be generalized to more realistic multisensory environments.

Given the online cross-sectional design of the current study, it provides evidence that the advantages associated with action video game experience do not extend beyond typical laboratory stimuli to more complex scenes incorporating some of the properties of real-world environments. As with many other studies, we cannot draw strong conclusions about the causal relationship between action video game experience and any effect (or lack thereof) from this study, as no training intervention was included in the protocol. Therefore, future work assessing performance within multisensory paradigms pre- and post-action video game training will be important to further evaluate the relation between action video game experience and cognitive abilities in real life.

The second aim of the present study was to assess whether there was a cross-modal advantage for AVGPs over NVGPs. By entering the normalized data into the analysis of performance on the search task in naturalistic scenarios, the results suggest that AVGPs do not benefit more from cross-modal cues than NVGPs (this conclusion can be interpreted within the statistical power of the present study, which is sensitive to medium to large effect sizes). These results are consistent with the previous study (Gao et al., 2018) which showed that AVGPs did not outperform NVGPs in their ability to integrate audiovisual stimuli. However, AVGPs display greater performance in some cross-modal tasks, such as simultaneity and temporal order judgments for visual and auditory stimuli (Donohue et al., 2010). Please note that in contrast to our task, TOJs require for effective segregation (rather than integration) of cross-modal events. This is because processing of the meaning of a complex naturalistic sound may require temporal integration due to the nature of the information (for a similar procedure, see Kvasova et al., 2019; and for a review see Vatakis & Spence, 2010). In the task used in our study, the sound was presented slightly earlier than the videos to capture attention and allowed us to study object-based congruency/incongruency effects in the integration of semantic information. Overall, this cross-modal semantic congruence benefited performance across the board, but it did so to the same extent in AVGPs and NVGPs. Overall, this pattern of results suggests that AVGPs do not benefit more than NVGPs from semantic audiovisual integration to direct their attention toward the target object. One explanation for this is that AVGPs and NVGPs may not necessarily employ different strategies or have developed different abilities when processing higher-level, semantic information used in the present study. Hence, the AVGP advantage consistently reported in other studies may be based on early, low-level processing. AVGPs have been shown to benefit from low-level spatial and temporal factors in their visual (Green & Bavelier, 2006; Greenfield et al., 1994; Schubert et al., 2015; Wong & Chang, 2018; Xuemin & Bin, 2010), auditory (Green et al., 2010; Stewart et al., 2019), or audiovisual (Zhang, Tang, Moore, & Amitay, 2017) search tasks. A recent meta-analysis confirmed a smaller impact on higher cognitive performance such as inhibition (*g =* 0.31, 95% CI [0.07, 0.56], *df* = 7.2, *p =* .02 and verbal cognition (*g =* 0.30, 95% CI [0.03, 0.56], *df =* 7.7 *p =* .033), while there was no significant impact on problem solving in cross-sectional studies (Bediou et al., 2018). Future studies need to explore whether extensive action video game experience has a positive impact on high-level cognitive functions or it is limited to low-level perception, spatial cognition, even if mediated by top-down attention.

From our exploratory analyses, we found that overall semantic congruence between sounds and target objects accelerated search latencies relative to neutral sounds, whereas semantic incongruence in the distractor-consistent condition did not produce, overall, a disadvantage relative to the baseline (neutral condition). Hence, consistent cross-modal cues benefit performance, while incongruent cross-modal cues do not hinder performance to a measurable extent. This result is consistent with other previous findings (Knoeferle et al., 2016; Kvasova et al., 2019; Kvasova, Spence, & Soto-Faraco., 2023; Iordanescu et al., 2010, 2011, 2013) and demonstrates that the semantic congruency effect can generalize beyond typical laboratory protocols and guide attention in a complex, multisensory environments. It also represent an extension of simpler, laboratory protocols (Iordanescu et al., 2008; Laurienti et al., 2003).

The design of this study allows us to calculate the false alarm rate and *d’* for the distractor-consistent condition which was a limitation of Kvasova et al.’s (2019) study. In line with the conclusions of that study, there was no difference in *d’* between distractor-consistent and neutral conditions. In the literature on cross-modal semantic congruence effects at large, the effects of target-consistent sounds are reliable, but the interfering effects of distractor-consistent sounds are often null or weak when found at all (Iordanescu et al., 2008; 2012; Knoeferle et al., 2012). This suggests that the cross-modal congruency effects can be more reliably interpreted as an advantage of congruent sounds, than distractor effects, which neither slowed down response time nor produced a more impulsive response than neutral conditions. An open question here is: If cross-modal semantics are clearly being processed, given the facilitation, why does semantic incongruence does not hinder performance? There are two possibilities for answering this question. One is that when visual stimuli are highly informative, a target-inconsistent sounds does not strongly distract participants from responding to the visual target. In this study, when the cue word is presented to participants, it activates a semantic network related to the target and creates an attentional template to that direct participants’ attention to other semantically congruent inputs from other modalities while inhibiting unrelated cross-modal information. In previous versions of the same task using the same materials, Kvasova et al (2019) included a ‘no sound’ condition. The results from this condition did not differ from the neutral condition (and for that matter, from the distractor-consistent condition). From these results, one could conclude that when sounds are incongruent or irrelevant with respect to the visual target, they may be not be processed sufficiently to produce a distractor effect. The second possibility is that in our study, some auditory stimuli may not be informative enough to direct participants’ attention and influence visual search because some of them are physically similar (e.g., the sound of the coin and keys) or semantically similar (e.g., the sound of musical instruments). Therefore, we cannot be sure that they were strong enough to play an effective distracting role above and beyond a neutral sound. To clarify this point, further studies should be conducted on the effect of high informative (hard to inhibit) versus low informative (easy to inhibit) cross-modal cues.

Finally, our findings can be somewhat informative to potential gender-based effects. The low prevalence of female AVGPs has typically led to a large gender imbalance in cross-sectional studies comparing AVGPs with NPVGs, which have been based almost exclusively on a male population (Cohen et al., 2007; Green & Bavelier, 2003; Green et al., 2010; Oei & Patterson, 2013). Here, we used a gender-balanced sample of participants, with equivalent numbers of males and females in both the AVPG and NVPG groups. Therefore, although sample size calculations were not focused on detecting possible gender effects, we believe that exploratory analyses using this variable in our pooled datasets (N=130; 49F, 51M) may provide a first impression, unavailable in previous studies. With regard to the putative general advantage of AVPGs in the search task as a whole (first hypothesis proposed), our results suggest that the AVGP advantage does not extend across the board. Playing AVGs was not associated with improved processing speed or attentional control capacity, and accuracy of responses (*d’* scores) remained unchanged across gender, as far as the sensitivity of our design allows us to tell.

Regarding the results of cross-modal advantage in male and female AVGPs and NVGPs, there were some gender variations. Please note that these variations must be interpreted in the context of an overall across-the-board crossmodal congruence benefit, without video-game experience group effects. Now, we observed that female AVGPs benefited more from cross-modal congruency in their RTs and were not distracted by cross-modal incongruency, whereas male AVGPs showed more distraction, with slower RTs on distractor-consistent trials. The increased processing speed observed in female AVGPs in a distractor-consistent condition can be viewed as more liberal behavior, in which female AVGPs respond faster while making errors equal to the neutral condition, compared to males who tended to trade time for accuracy (see Figure 5C and 5D). These findings suggest a potential difference in response strategy between experienced male and female AVGPs when dealing with cross-modal cues, to be confirmed in the future. With due caution, this finding may be consistent with some previous studies suggesting that there are gender differences in selective attention and spatial cognition (Evans & Hampson, 2015; Halpern, 2013; Merritt et al., 2007; Posner & Marin, 2016; Stoet, 2017). Another study investigated the gender-specific effects of AVGP experience and showed that playing AVGs reduced the differences in spatial cognition between male and female (Feng, Spence, & Pratt, 2007). Further studies should be performed to follow up these findings. For now, one conclusion that can be drawn from this study is that there may be potential for studying gender differences in the effects of AVGP, an area that has been neglected in all studies to date.

**Figure 5.**
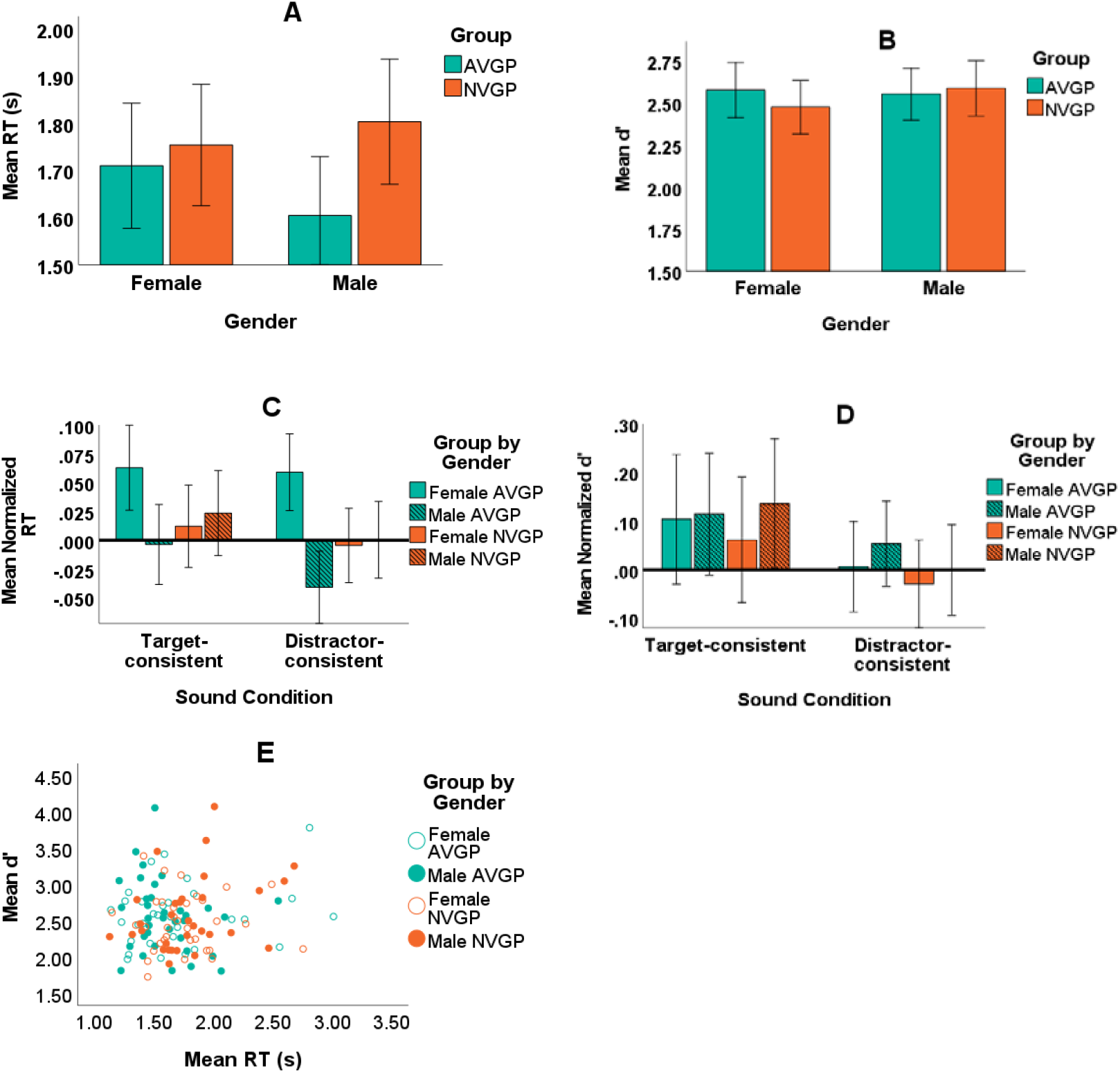
**(A)** Visual search Mean reaction times for correct responses to target-present trials, plotted separately for male and female AVGPs and NVGPs. (B) Total visual search accuracy (*d’*) plotted for male and female AVGPs and NVGPs. (C). Normalized reaction times in the target-consistent and distractor-consistent conditions plotted for male and female AVGPs and NVGPs. (D) *D’* values of the three sound conditions plotted separately for male and female AVGPs and NVGPs. (E) Individual *d’* scores plotted against average RTs pooled across the four group*gender. In all plots, error bars indicate the standard error of the mean.

It is worth mentioning that this study was performed online, with the experiment presented in a browser on the participants’ personal device (tabletop computer or laptop). This contributes to the generalization of our findings to more ecological situations. Despite our efforts (see Methods), the luminance of the auditory stimuli and video clips may have varied between participants. We assume that this variation was similar between the groups (AVGPs vs. NVGPs vs. male vs. female). There is reason to assume that such group differences were negligible, as our data replicated the finding of the cross-modal semantic effect, which is very similar to the laboratory-based study by Kvasova et al. (2019).

Another potential limitation of the study is the possibility that home setups may account for the reaction time differences observed in Experiment 1. It seems plausible that AVGPs performed the experiment using a different computer setup than NVGPs: perhaps they had ergonomic keyboards or used external monitors (as opposed to laptops). They may also have had more comfortable (gaming) chairs and a dedicated room with fewer distractions. Although research has shown that there are no significant differences for RTs in lab-versus web-based tasks (Hilbig, 2016), hardware can cause potential delays in time measurement that ranging from 0.0006 to 80 ms for keyboards and 0.0005 to 68 ms for monitors (Crocetta and Andrade, 2015). However, further work in other natural contexts (e.g., virtual reality or other situations that do not involve sitting in front of a display) is needed to evaluate the extent to which the observed advantages can be generalized.

## Conclusions

The results of the present study suggest that the AVGP advantage in attentional tasks does not extend to complex search tasks with naturalistic, multisensory scenes. We also failed to demonstrate that AVGP experience confers a specific advantage in exploiting cross-modal cues, or in resisting distractors, compared to NVGPs. Our data, however, shed some light on semantic aspects of multisensory integration by generalizing (and confirming) previous laboratory findings on the semantic congruency/incongruency effect on cross-modal interactions to an online setting. Other findings from exploratory analyses with gender suggest that there may be strategic differences in how this advantage is achieved. In order to gain a full understanding of the AVG advantage in males and females, it will be important to include female participants in this area of research and to compare results between genders more systematically.

## DATA AVAILABILITY STATEMENT

The datasets generated for this study are available at https://osf.io/sgy5a/.

