## Supplementary materials for "Addressing the Association Between Action Video Game Playing Experience and Visual Search in Naturalistic Multisensory scenes"

Submitted to: *Multisensory Research*

**Supplementary Materials**

**Experiment 1**

We ran pre-registered analyses for the first hypothesis with all 85 participants (the included and all excluded ones for the accuracy criteria) on both RT and accuracy. The results of a one-sided t-test on mean RTs showed significant differences between groups(t_(83)_ = 2.30, *p* < 0.05, *Cohen’s d* = 0.50, BF_10_ = 4.33), and significant differences between groups on *d’* total (t_(83)_ = 3.06, *p* < 0.05, *Cohen’s d* = 0.4, BF_10_ = 1.90) with a small effect size and anecdotal evidence for H_1_. As can be seen in Figure S1, there are some outliers in both graphs, especially in the distribution of d’ in the NVGP group. Exclusion criteria helped us to filter outliers and minimize ambiguity in the data.

**A**

*

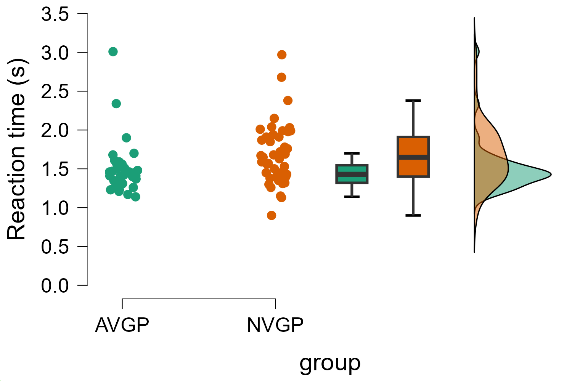

##### B

*

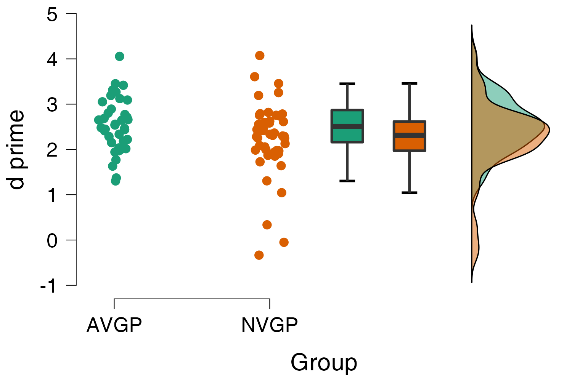

**Figure S1. (A)** Visual search average reaction times toward targets and error rates plotted separately for Action Video Game Players (AVGP) and Non-Video Game Players (NVGP). **(B)** Visual search accuracy (d’) was plotted for the two groups. Significant differences are indicated by asterisks (*p-value<0.05).

**Experiment 2**

We repeated the above analyses for the first hypothesis with all 96 data from Experiment 2 (the included and all excluded ones for the accuracy criteria) on both RT and accuracy. The results showed that there isn’t any significant difference between the two groups on RTs (t_(94)_ = 1.18, *p* = 0.12, *Cohen’s d* = 0.50, BF_10_ = 4.33), as well as d’ scores on *d’* total (t_(94)_ = 0.33, *p* = 0.37, *Cohen’s d* = 0.4, BF_10_ = 1.90). These results are inconsistent with the results of Experiment 1 and showed that even without exclusion criteria the differences between the two groups were so small to be detected by this experiment. Figure S2 depicts the distribution of the data for RTs and d’ scores. It is clear that the exclusion criteria helped us to filter outliers and minimize ambiguity in the data.

**A**

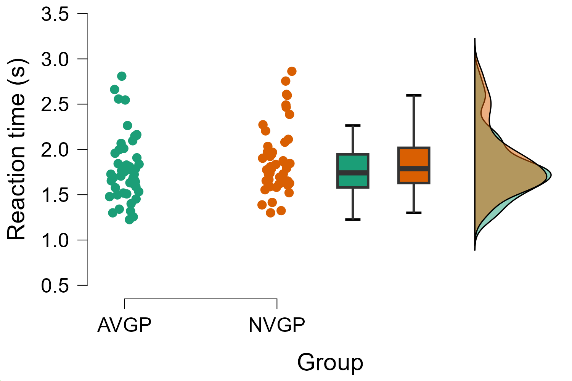

##### B

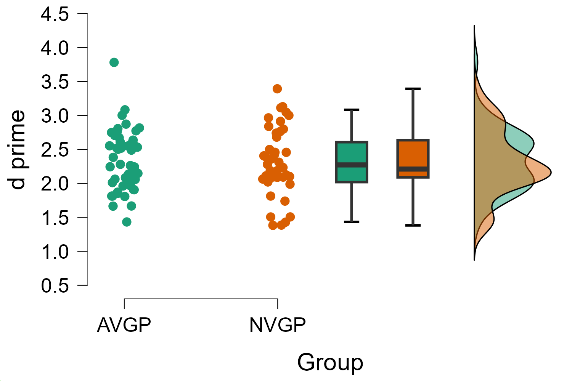

**Figure S2. (A)** Visual search average reaction times toward targets and error rates plotted separately for Action Video Game Players (AVGP) and Non-Video Game Players (NVGP). **(B)** Visual search accuracy (d’) was plotted for the two groups.
